# Image-based taxonomic classification of bulk biodiversity samples using deep learning and domain adaptation

**DOI:** 10.1101/2021.12.22.473797

**Authors:** Tomochika Fujisawa, Víctor Noguerales, Emmanouil Meramveliotakis, Anna Papadopoulou, Alfried P. Vogler

## Abstract

Complex bulk samples of invertebrates from biodiversity surveys present a great challenge for taxonomic identification, especially if obtained from unexplored ecosystems. High-throughput imaging combined with machine learning for rapid classification could overcome this bottleneck. Developing such procedures requires that taxonomic labels from an existing source data set are used for model training and prediction of an unknown target sample. Yet the feasibility of transfer learning for the classification of unknown samples remains to be tested. Here, we assess the efficiency of deep learning and domain transfer algorithms for family-level classification of below-ground bulk samples of Coleoptera from understudied forests of Cyprus. We trained neural network models with images from local surveys versus global databases of above-ground samples from tropical forests and evaluated how prediction accuracy was affected by: (a) the quality and resolution of images, (b) the size and complexity of the training set and (c) the transferability of identifications across very disparate source-target pairs that do not share any species or genera. Within-dataset classification accuracy reached 98% and depended on the number and quality of training images and on dataset complexity. The accuracy of between-datasets predictions was reduced to a maximum of 82% and depended greatly on the standardisation of the imaging procedure. When the source and target images were of similar quality and resolution, albeit from different faunas, the reduction of accuracy was minimal. Application of algorithms for domain adaptation significantly improved the prediction performance of models trained by non-standardised, low-quality images. Our findings demonstrate that existing databases can be used to train models and successfully classify images from unexplored biota, when the imaging conditions and classification algorithms are carefully considered. Also, our results provide guidelines for data acquisition and algorithmic development for high-throughput image-based biodiversity surveys.

## INTRODUCTION

Biological identifications increasingly rely on machine learning algorithms that use photographic images to place unidentified specimens into a taxonomic classification. As these methods are proving to be very powerful especially for identification of the species-rich and morphologically diverse insects, it is now possible to place a specimen with high confidence against curated image libraries, *e.g*. those obtained from pinned museum collections (Buschbacher *et al*. 2020; Hansen *et al*. 2020). With the rapid increase of such images, machine learning can greatly increase the capacity for species identification without putting demand on scarce taxonomy experts (Valan *et al*. 2019; Høye *et al*. 2021). The methodology therefore is likely to play a major role in the taxonomic endeavour in future, and potentially deep learning can have similar impacts on the practice of taxonomy as the revolution of DNA barcoding and metabarcoding some 20 years ago, or it could work in concert with these molecular approaches (Høye *et al*. 2021; Wührl *et al*. 2021; Yang *et al*. 2021). However, the true potential and possible limitations of algorithmic methods for exploiting the information contained in specimen images remain to be established, as the various applications and choice of machine learning algorithms continue to be refined (Valan *et al*. 2019; Romero *et al*. 2020).

The greatest challenge for modern taxonomy probably is the study of highly diverse and poorly studied biota and geographic regions, harbouring many undescribed species (Costello *et al*. 2013). In particular, in studies of invertebrate diversity, such as those from tropical forest canopy or the soil, huge numbers of specimens are collected and subsequently need to be classified and counted as part of ecological and environmental studies (Novotny *et al*. 2007; Caruso *et al*. 2018). Imaging of these specimens is comparatively fast with the help of recently described automated imagers (Ärje *et al*. 2020; Wührl *et al*. 2021) or by taking high resolution images of large sets of specimens in a single photo, which can then be cropped to represent single individuals for subsequent classification (Hudson *et al*. 2015; *https://www.site100.org*). Automated classification based on these images would remove the need for manual identification by taxonomic experts who individually can handle only a small portion of the diversity spectrum usually encountered in such studies (Basset *et al*. 2012), and thus may help to provide rapid assessment of threatened arthropod assemblages, where speed is a priority.

In machine learning, images are classified against a class of defined objects, *e.g*. the images of a particular taxonomic group. A model trained to separate the types of images in this source is used to classify unlabeled objects in the target, such as an unknown set of specimens in a sample. Most recent studies used convolutional neural networks (CNN, LeCun *et al*. 2015) for the task of image classification. Because of the lack of images for training the full parameters of a CNN model, approaches like fine-tuning of the existing CNN (Ärje *et al*. 2020) or feature transfers from the pre-trained CNN (Valan *et al*. 2019) are commonly used in biodiversity studies, following the successful applications of pre-trained CNN outputs as generic image features (Donahue *et al*. 2013; Razavian *et al*. 2014). These methods of transfer learning have already shown great power in taxon annotation and detections applied to insect specimens, and in some cases surpass the capabilities of trained taxonomists (Valan *et al*. 2021).

Yet, applications of image classification algorithms for insect biodiversity research have mainly been limited to narrow tasks and specific target sets, such as pinned museum specimens (Valan *et al*. 2019; Hansen *et al*. 2020), aligned body parts (Buschbacher *et al*. 2020; Klasen *et al*. 2021), or small target groups of a few species (Ärje *et al*. 2020). In most of these studies the unlabeled (target) set is from the same category, *i.e*. the target taxa at species or higher hierarchical levels are included in the training set. However, as collecting labeled datasets is the most laborious part of machine learning applications, it is desirable that models trained in one dataset can be used for prediction tasks of others (called ‘domain adaptation’, *e.g*. Kouw & Loog 2021). While methods for domain adaptation have been successfully applied to fields such as medical image classification (Guan & Liu 2021), they have not been used in the context of biodiversity surveys. Their application would be particularly useful in classification of specimens from complex, mixed trap samples, especially from unexplored areas whose components are unlikely to be present in the training set.

Building such an image-based classification system may be complicated by several factors because the assumption of identical distributions between the source and target datasets rarely holds. Capture bias is a well established problem in machine learning, as objects appear in different contexts (location, lighting, background, etc.) or are taken on different imaging devices. Images of arthropods may be from collection specimens taken in a fairly standardised position and lighting conditions (Valan *et al*. 2019; Hansen *et al*. 2020), or may be obtained directly from bulk samples and photographed either singly (Raitoharju *et al*. 2018; Valan *et al*. 2019; Wuhrl *et al*. 2021) or cropped from large-field composite images (Buschbacher *et al*. 2020; Hansen *et al*. 2020). Images thus feature different aspects of the specimens and differ in illumination and magnification, which affects the recognition of key features (Raitoharju *et al*. 2018; Ärje *et al*. 2020). Other issues are unrelated to differences in image acquisition, but result from the biases of defining the semantic categories (Tommasi *et al*. 2017). Such “category bias” may arise from inconsistent labeling, either due to the application of different taxon concepts used for naming of species and higher taxa, or due to specimen misidentification. The resulting noisy or incorrect data labels then reduce the effectiveness of the model. In addition, in particular in higher taxonomic categories, the same name is assigned to visually different images due to the distributional shift of subclasses (*e.g*. different genera representing a family in the source and target). Furthermore, in general cross-dataset applications, the model can encounter a category which is missing in the source training data, *e.g*. a new family may be present. The treatment of such anomalous (or *out-of-distribution*; see Tabak *et al*. 2018) samples affects the reliability of the biodiversity assessment. As more variation is encountered, to fully learn the structure of the data, the model should scale with the size and complexity of the training data.

In practice, due to these problems of intra-class variability and the inconsistencies of the photographs, the success of deep learning in taxonomy to date has been in situations where a bespoke image library is available that holds a narrow representation of the query taxa and images under the same aspect and imaging conditions (brightness, angle, magnification, *etc*; Buschbacher *et al*. 2020; Valan *et al*. 2021). However, the utility of these methods remains largely untested in the application to samples from poorly characterised species, as those from bulk arthropod sampling of fauna in unexplored areas that have not been encountered previously in the image reference set. Ideally, such samples would be identifiable against images drawn from other sources, for example an image database of well characterised regional communities and taxa obtained elsewhere.

Here, we test the usage of deep learning for insect classification based on bulk-sample images in real-world scenarios of high-throughput biodiversity surveys. We characterise bulk samples of poorly known communities of Coleoptera (beetles), where high species diversity, idiosyncratic morphological variation, and the constraints of bulk images provide a challenging, but realistic situation for the machine-based classification. The aim was a classification at family-level, as the first step towards an in-depth study of biodiverse regions and complex ecosystems around the globe. Specifically, we address three parameters to affect the prediction accuracy: (i) the quality of images, especially the resolution of the image using standard macrophotography versus high-resolution stacking technology; (ii) the transferability of identifications across communities from different habitats and continents, *i.e*. when the input subclass is not present in the training data; and (iii) the impact of the size and complexity of the training set. We conclude that the prediction success for the classification depends on the beetle family (*i.e*. some classes are more easily predicted) and the number of images and the image acquisition methods of the training set.

## MATERIALS AND METHODS

### Sample collection and taxon selection

As the target for classification, we used a collection of leaf-litter samples from a total of 46 sites distributed across five forest habitats of the Troodos mountain range of Cyprus (Fig. 1). These samples were processed as described by Noguerales *et al*. (2021) to extract bulk Coleoptera specimens from the substrate using a Berlese apparatus. During sample processing, single specimens were selected from the bulk samples and individually processed. Bulk samples and single specimens were preserved in 100% ethanol and subsequently photographed following two different imaging protocols (see below). For more details on soil sampling and habitat descriptions, see Arribas *et al*. (2016) and Noguerales *et al*. (2021), respectively. During sample processing and imaging, the most common families/subfamilies, with 5 or more photographs per taxonomic rank, were identified and used for downstream analysis. The chosen families were: Brentidae, Carabidae, Chrysomelidae, Cryptophagidae, Curculionidae, Latridiidae, Leiodidae, Melyridae, Ptilidae, Staphylinidae:Scaphidiinae, Staphylinidae (excluding Scaphidiinae) and Tenebrionidae (Table S1).

**Figure 1.**
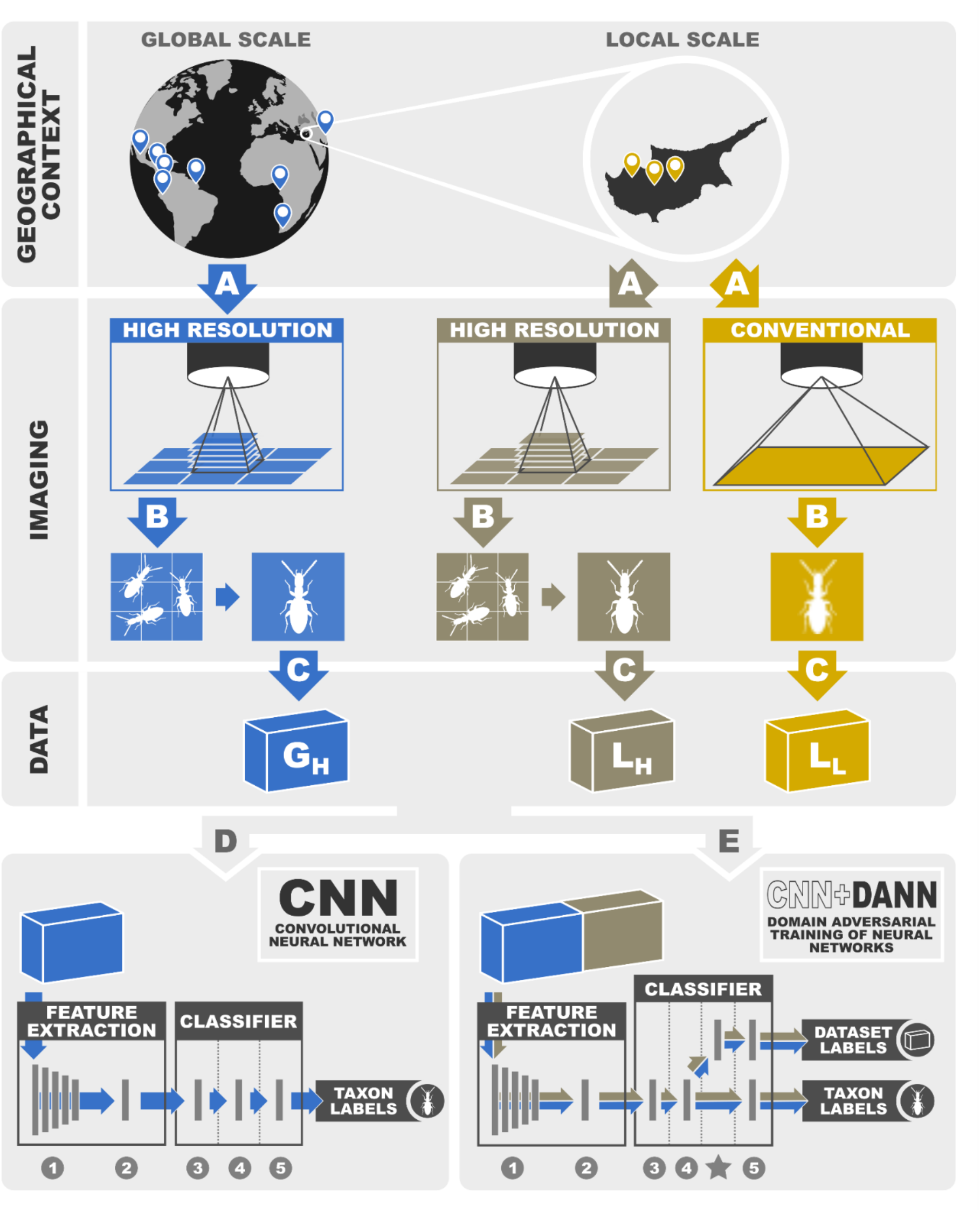
Schematic diagrams summarizing the experimental workflow of the study, depicting the geographical context and the different imaging procedures for generating the three image datasets: G_H_, Global High Quality, L_H_, Local High Quality; L_L_, Local Low Quality. Taxa classification was performed using two alternative deep learning algorithms: convolutional neural network (CNN) and domain adversarial neural network (DANN). For more details on algorithm-specific architectures, see Figure S1.

### Image data acquisition

#### Local High Quality (L_H_) and Local Low Quality (L_L_) datasets

Specimens from bulk-samples were air-dried and placed at regular distances onto filter paper in a Petri dish. In cases of large disparity in body size, we split the bulk samples into different size categories which were separately photographed in order to improve the focus and resolution across all specimens regardless of their body size. As much as possible, specimens were positioned for photography in the dorsal view.

Bulk-sample photographs were taken using a Zeiss AXIO Zoom.V16 Stereo Zoom Microscope equipped with a Zeiss AxioCam HRc (High Resolution 13 Megapixels Color Microscope) camera at the Imaging and Analysis Centre at the Natural History Museum (NHM) in London, United Kingdom. This on-axis instrument with motorised focus drive and motorised stage enables large high-resolution images by dividing the field into regular tile-images which were subsequently *xyz* stitched. Depending on the sample size, photographs were taken by dividing them into 16-64 tiles, each with 25-30 slices (*z-*stacks) using the Zeiss NEO 2 Blue Edition software. We rendered *z*-stack images with the Helicon Focus v.5.3.14 software (*https://www.heliconsoft.com*) using the pyramid-based algorithm (‘Method C’) and default parameters. Focus stacking was also performed using the depth-map algorithm (‘Method B’) in Helicon Focus with a radius value of 8 and a smoothing parameter of 4, yielding qualitatively similar images to the former method. Consequently, only photos from ‘Method C’ were used for downstream analyses.

Finally, we manually cropped single specimen photos from the bulk-sample images using INSELECT v.0.1.35 software (Hudson *et al*. 2015). After some minor corrections of bounding edges, cropped single-specimen images were exported and taxonomically identified at the family/subfamily level by the authors. Only whole-bodied specimens were considered for further analyses. The cropped images were resized to 255×255 pixels for subsequent classification tasks. When an image was not an exact square, the edges were padded using the average pixel value of the outermost portions of the image to enforce a square shape.

The individual frames cropped from the bulk samples were denoted the *Local High Quality* (L_H_) data set, referring to the fact that they were obtained from a local area and thus represent a small taxonomically confined set, and taken at high image resolution. The L_H_ dataset represented the best case scenario, where high-resolution training images of local samples are obtained under controlled conditions with high-performance imaging equipment.

During the processing of local bulk samples, the selected individual specimens were also photographed using a conventional stereoscope NIKON SMZ1270i equipped with a NIKON DS-Fi3 Microscope Camera (5.9 megapixels) controlled by the NIKON DS-L4 v.1.5.0.3 control unit. These photographs were intended to represent a more realistic scenario of local specimens being photographed during field sampling and sample sorting in local lab facilities using conventional instruments. These images were denoted *Local Low Quality* (L_L_) dataset.

#### Global High Quality (G_H_) dataset

We also obtained a wider sample of images from a global catalogue of Coleoptera specimens available at *https://www.flickr.com/photos/site-100/.* These images had been obtained from local sampling campaigns at 11 sites throughout Central America, Africa and Southeastern Asia (see Table S2) and photographed in bulk using the Zeiss AXIO Zoom, as described above, while others were individually taken at high-resolution on a single lens reflex (SLR) camera (Canon EOS 500D) and macro lens (Canon MP-E 65mm f/2.8 1-5x Macro). Helicon Focus software was used to render *z*-stack images, as aforementioned described. This dataset was denoted the *Global High Quality* (*G_H_*) dataset. For each of the selected families, all specimen photographs available for the respective sites were used. Relative numbers of available specimens per family were usually correlated across sites, with greatest numbers in Staphylinidae. The numbers of images in the three data sets are shown in Table S1.

### Image classification with neural network (NN)

#### Feature transfer and neural network classifier

We employed the strategy of feature transfer from the pre-trained convolutional neural network (CNN) proposed by Valan *et al*. (2019). We chose the outputs of the fifth convolutional block of the VGG19 model after 2-dimensional average pooling as a set of features for an image, based on the results of Valan *et al*. (2019) and our pilot analyses. These 512-dimensional image features were used for the classification with a neural network. Alternative pre-trained models, VGG16 and ResNet, were also tested, but were outperformed by the VGC19 model.

The neural network classifier consisted of two fully connected (FC) layers with ReLU activation and a softmax output layer (Fig. 1 and Fig. S1). The dropout was applied after the FC layers with a dropout rate of 0.6. The neural network was trained with the stochastic gradient descent algorithm with the softmax cross-entropy loss. The numbers of units in the two FC layers (512 and 256 for the first and second FC layers respectively) and the dropout rate were determined by five-fold cross-validation with a random 200 images of the G_H_ dataset, and these hyperparameters were used throughout all classification tasks in this study.

#### Within-dataset classification

The CNN model was trained with *N* images randomly selected from the dataset and predicted the class of *n* test images randomly selected from the rest. *N* ranged between 100 and 700 for L_H_, 50 and 250 for L_L_, and 100 and 900 for G_H_. The number of test images, *n*, was set to 200 for L_H_ and G_H_, and 50 for L_L_ due to the small size of the dataset. To evaluate the consistency of prediction accuracy, ten replicates were generated for each scenario of *N* images. The output of the final softmax layer was used as the prediction probability of each class, and the image was classified to the class with the highest prediction probability. The accuracy of the prediction was measured as the proportion of successful predictions in the test set, 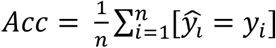, where 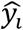 is the predicted class of the *i*-th image, *y_i_*, the true class, and 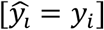 is 1 if 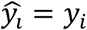 and 0 otherwise.

The classification performance for each class was measured by the multiclass recall rate, multiclass precision and the F1-score. Recall rate of class *c* is defined as a proportion of correct predictions of *c* out of the actual number of images of *c*, 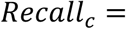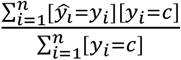. Multiclass precision is defined as a proportion of correct predictions of *c* out of the number of images predicted as c, 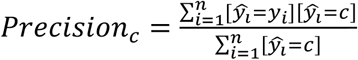. The F1-score is a harmonic mean of the multiclass recall rate and precision. Thus, while the recall rate is interpreted as the fraction of images of a class present in the sample that are selected, the precision quantifies the fraction of the images predicted as members of a class that are actually correct. The F1-score represents the overall performance of a classifier with respect to these two measures.

For the L_H_ dataset, we tested the effects of a locality-wise sampling strategy on prediction accuracy. Instead of pooling all images and randomly selecting a training set from them, we split the images based on their sampling sites within Cyprus by generating separate training sets composed of images from *S* randomly selected sites in the Troodos mountains. Then, the NN predicted 200 images from the sampling sites not present in the training set. The number of sites, *S*, ranged from 4 to 28 with an interval of 4.

In addition, we used an L_H_-trained model to predict the class of 16 high-quality images belonging to 8 families/subfamilies, Coccinellidae, Elateridae, Endomychidae, Hydrophilidae, Laemophloeidae, Phalacridae, Scarabaeidae and Scydmaeninae, which were not present in the training data, and thus test the effect of unknown inputs (hereafter *out-of-distribution* samples) on the classification (see Tabak *et al*. 2018).

#### Between-datasets classification

For the between-dataset prediction the CNN model was trained with a source dataset to predict images from a different target dataset. The NN was trained with *N* images randomly selected from the source dataset, which was then used to predict all images of the target dataset. The target accuracy, *Acc_T_* was measured as the proportion of successful predictions of the target images. The baseline accuracy within the source dataset, *Acc_S_*, measured in the within-dataset classification was compared with the target accuracy, *Acc_T_*. The accuracy reduction, Δ*Acc(S, T)* = *Acc_S_* − *Acc_T_*, was recorded as a measure of transferability between the datasets. High Δ*Acc* indicates large reduction of accuracy, hence difficulty in transfer. We ran the above procedures for three source-target pairs (training dataset → predicted dataset), G_H_ → L_H_, G_H_ → L_L_ and L_L_ → L_H_. These settings simulate two alternative scenarios: (i) a global image database is used to predict local samples (G_H_→L_H_ and G_H_→L_L_) and (ii) conventional images, as those representing single-specimen photographs by local taxonomists, are used to predict local high-resolution images (L_L_→L_H_).

Divergence between the source and target datasets was measured with a dataset classification error. A linear support vector machine (SVM) was trained to classify images to the source or target dataset with the features of 200 randomly selected images from both datasets. Conversely to above analyses, here the model was trained to classify datasets instead of taxa. Then, a classification error of the SVM, *ε_source-target_*, was measured as a proportion of incorrect predictions of 200 test images sampled from the two datasets. An intuitive interpretation of this measure is that the dataset classification task is harder when the feature distributions between two datasets are more similar. Therefore, a large classification error indicates high similarity between source and target datasets. This approach is commonly used to measure the dataset bias (Tommasi *et al*. 2017).

#### Between-datasets classification with domain adversarial training

In addition to the ordinal CNN setups described above, we employed the domain adversarial training of neural networks (DANN, Ganin *et al*. 2016) to improve the accuracy of the between-dataset classifications. DANN uses labeled images from the source as well as unlabeled images from the target dataset in its training process to improve the target predictions. The DANN model jointly predicts the taxon (class label) of the source images and the dataset (domain) of all input images (as in the previous section) by adding layers for the dataset classification to the classifier (Fig. S1). The training procedure then optimizes the model parameters in the shared part of the network to not only minimize the loss of the label classifier (taxon prediction) but at the same time to *maximize* the loss of the domain classifier (dataset prediction). This adversarial training procedure optimizes shared intermediate features to be invariant between the two domains, and hence the model can generalise across them, which potentially improves the accuracy in target predictions. In this study, a softmax layer with binary cross entropy loss was added as a dataset classifier to the NN after the second FC layer. The regularization parameter, λ, which controls the relative importance of the two classifiers, were set to λ = 0.1, 0.5 and 1.0, and the best performing results (λ = 0.1) were reported.

The performance of the DANN method was measured with procedures similar to those in the previous section. A mixed set of images of size *N* was randomly selected from target and source datasets, and training was done using taxon labels from the source images and dataset labels for all images. Next, 400 mixed test images were predicted, and their source and target accuracies and their difference were recorded. We applied the DANN to the three pairs from the previous section. The total number of images *N* ranged between 300 and 800 for L_L_→L_H_, 400 and 1400 for G_H_→L_H_, and 300 and 1000 for G_H_→L_L_. The proportions of source images were 0.3, 0.67 and 0.83 for L_L_ →L_H_, G_H_ →L_H_ and G_H_ →L_L_ respectively, which yielded training images from the source similar in number to the other training setups. The effect of DANN on target accuracy was tested using linear regression with the model type and the number of images as explanatory variables. Models of neural networks were implemented in Python with Keras (*https://keras.io*) and TensorFlow (*https://www.tensorflow.org*) libraries, and all statistical analyses were conducted with R (R Core Team 2021).

## RESULTS

### Performance of within-dataset classification

#### Effects of datasets and the number of images

The accuracy of within-dataset classification and the effect of the number of training images varied among datasets. The accuracy for the L_H_ samples generally improved with an increasing number of training images and reached an average of 96% with 700 images (Fig. 2a). The maximum classification accuracy for the L_H_ was 98%. When the locality-wise training was performed, the accuracy slightly decreased to an average of 91% with 28 localities, which were roughly equivalent to 680 images.

**Figure 2.**
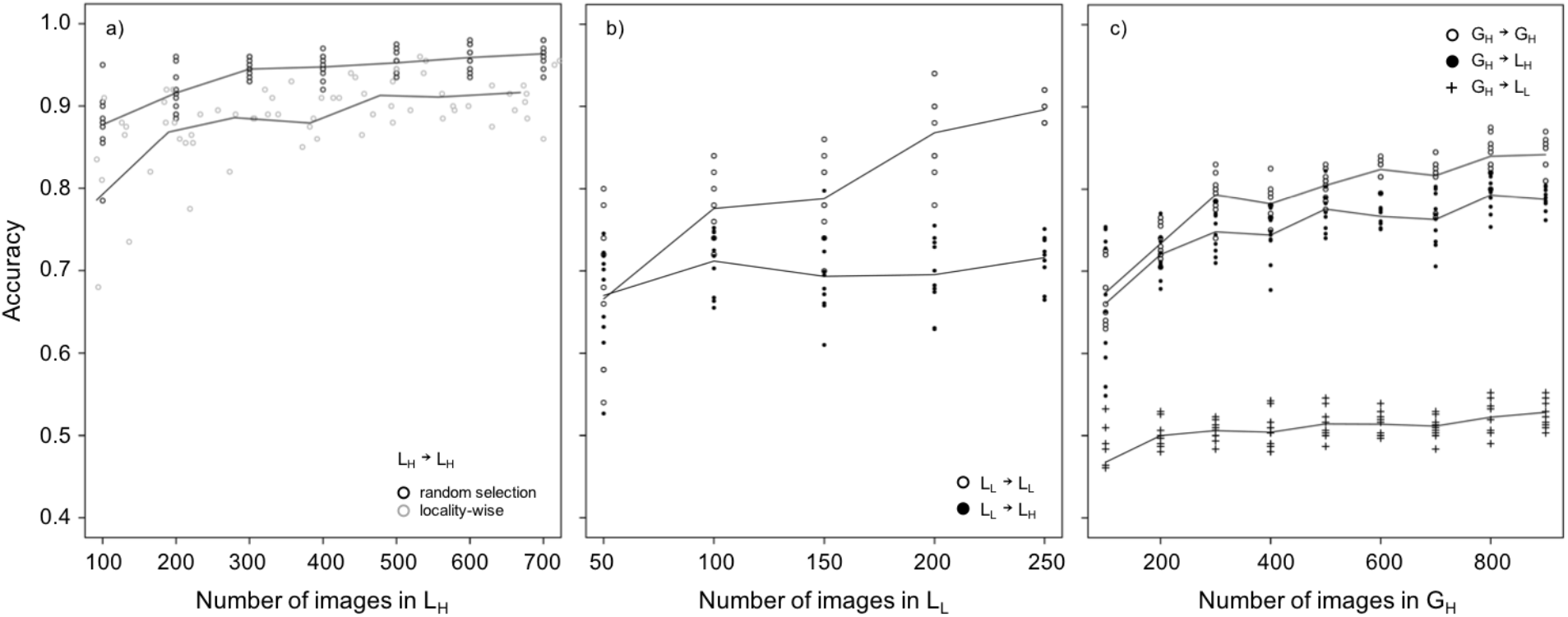
Effect of increasing the number of images on prediction accuracy. Training the convolutional neural network (CNN) on a subset of images and prediction of the class of images for (a) Local High Quality (L_H_) images selected either at random or under locality-wise selection for predicting the class of L_H_ images, (b) Local Low Quality (L_L_) images for training and predicting the class of either L_L_ or L_H_ images, and (c) Global High Quality (G_H_) images for training and predicting the class of either L_L_, L_H_ or G_H_ images.

The within-dataset classification accuracy of the L_L_ images, taken by a conventional stereoscope and camera, was generally lower compared to the L_H_ dataset. The accuracy increased monotonically with the increasing number of images and reached an average of 89% with 250 images (Fig. 2b). The within-dataset classification accuracy of the G_H_ images was also lower compared to the L_H_. The improvement of accuracy was slower than for the other datasets, and the average accuracy was 84% with the maximum number of 900 images (Fig. 2c).

Classification error was visualised as a scaled confusion matrix for a trial with 400 training images in the L_H_ random sampling (Table S3). The large taxonomic groups were correctly classified in most cases. For example, four families (Carabidae, Curculionidae, Ptiliidae and Staphylinidae) were classified with more than 95% recall rate (Fig. 3a), while the remaining taxa had widely different recall rates ranging from 0% to 82% (Fig. 3a). In the extreme case of the family Melyridae, with the lowest number of available images (*n* = 5), no images were predicted correctly (Fig. 3a). When a taxon had >50 images, its recall rate and precision approached 1.0 (Fig. 3a,c). The F1-scores showed a similar pattern, *i.e*. for those images that were called to be members of a taxon, these predictions were generally correct (Fig. 3e). Class-wise recall rates and F1-scores showed a strong positive correlation with the number of images (rho = 0.81 and 0.85, respectively; Fig 3a,e). The effect of the number of images on the class-wise precision was also positive, but slightly weaker (rho = 0.41, Fig. 3c). Failed predictions included ventral views of insect bodies, specimens with missing body parts or multiple specimens in a single image (see Fig. S2). Apart from these irregular images, most failed predictions were for taxa represented by <20 images (Fig. 3a,c,e).

**Figure 3.**
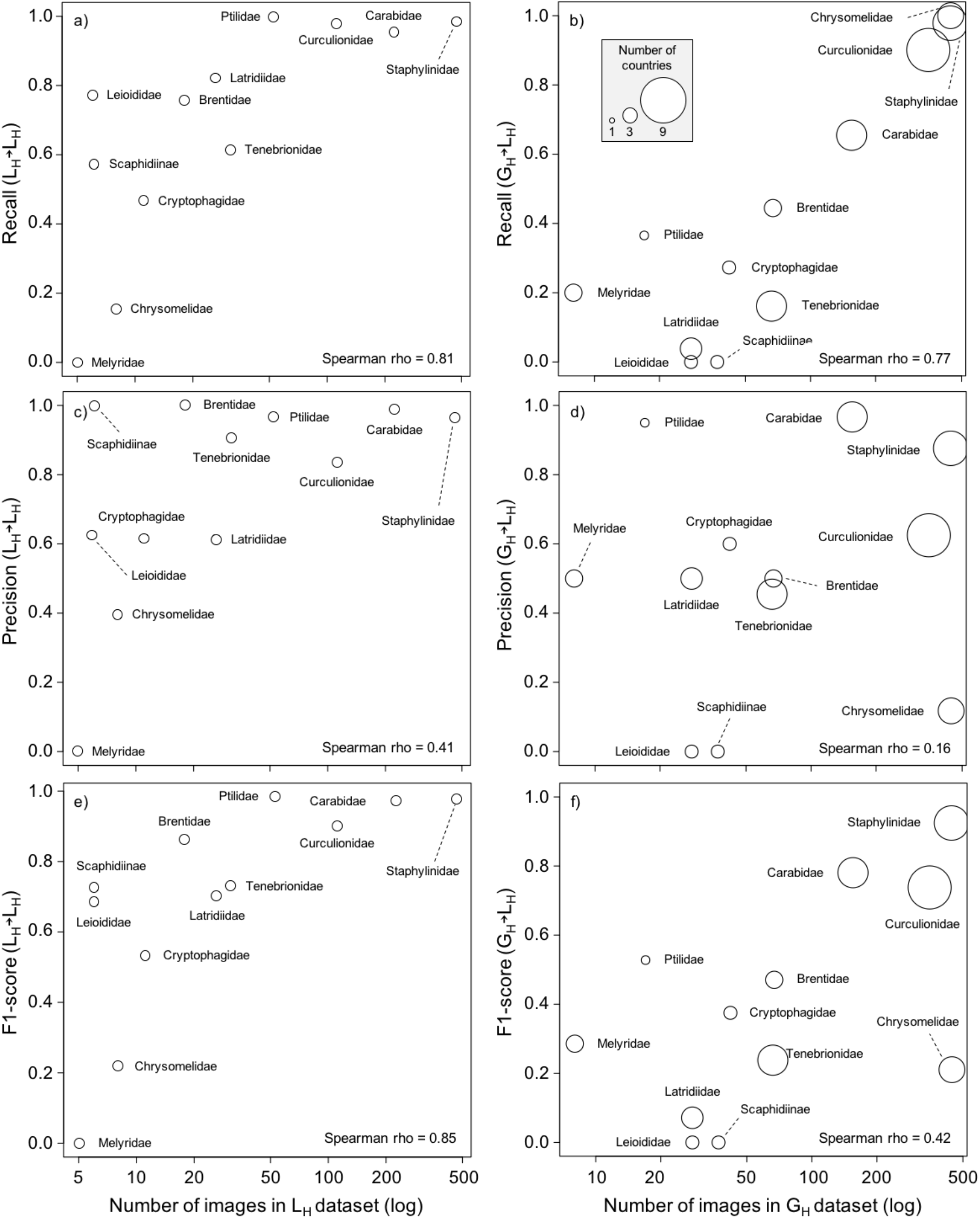
Effect of the increasing number of images on recall rates (panels a and b), multiclass precision (panels c and d) and F1-scores (panels e and f). We used 400 randomly selected Local High Quality (L_H_) images for training and predicting the class of L_H_ images (within-dataset classification) (left panels), and 800 randomly selected Global High Quality (G_H_) images for training and predicting the class of L_H_ images (right panels). Note that x-axes representing the number of images on a logarithmic scale. Circle sizes represent the number of countries where samples of a given family were collected from (as a proxy of intra-family morphological variation).

#### Prediction probabilities and out-of-distribution samples

Prediction probabilities for the successful predictions (average 0.98) were overall higher than for the failed predictions (average 0.79, Fig. 4), when using the L_H_ dataset with 400 training images. For images assigned to families not present in the training data (that is, *out-of-distribution* samples), the prediction probabilities were also lower on average than for the successful predictions (average 0.83, Fig. 4). However, four samples were predicted incorrectly with probabilities of more than 0.95 (Fig. 4). For example, three images of Coccinellidae, Hydrophilidae and Phalacridae were classified as Ptiliidae with probabilities >0.95.

**Figure 4.**
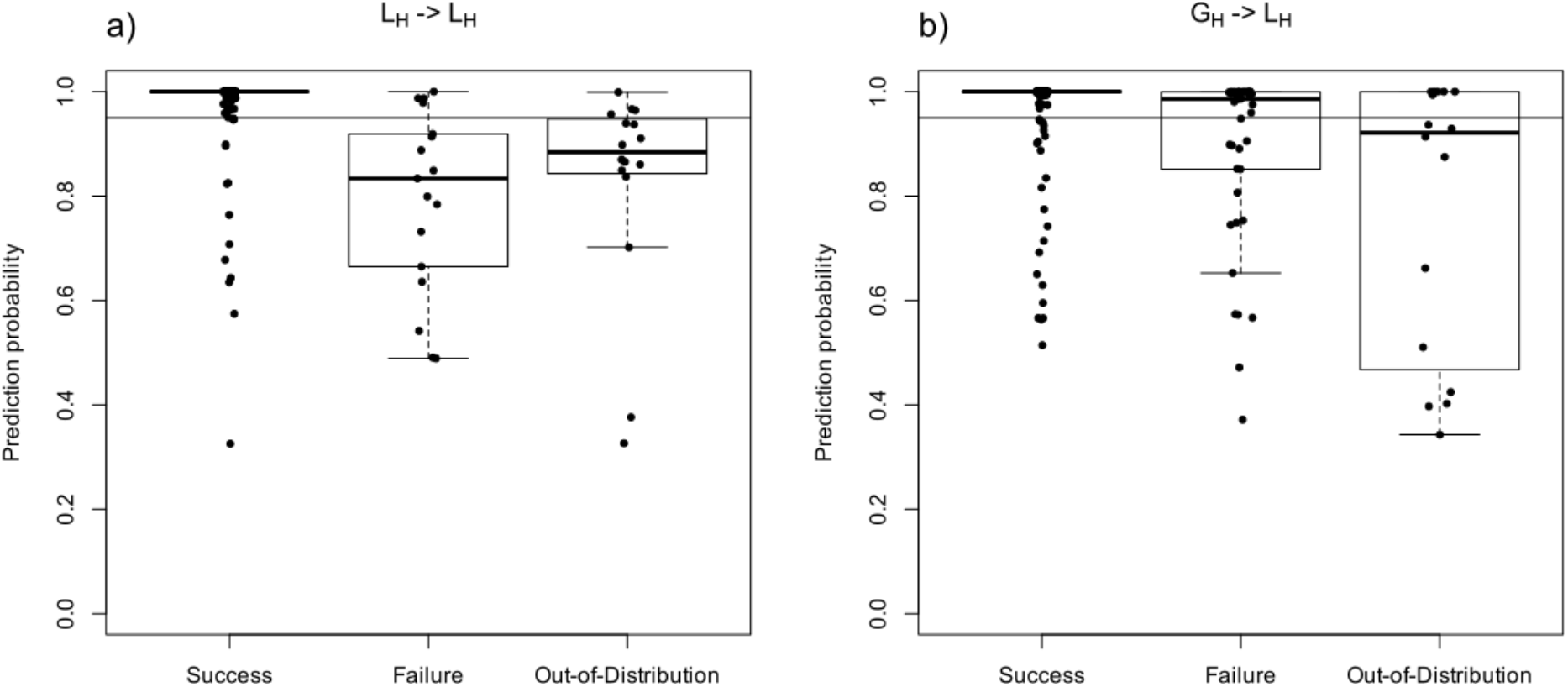
Prediction probabilities for the successful, failed and out-of-distribution predictions at a 0.95 threshold (horizontal line). (a) Intra-dataset predictions of L_H_ images using 400 randomly selected images for training. (b) predictions of L_H_ images using 800 G_H_ images for training.

To detect the failed predictions, we set conservative threshold values for the prediction probabilities and marked samples below the threshold as potential misclassification. When the threshold value was set to 0.95, 92% of successful predictions were retained while 76% of failures and 75% of *out-of-distribution* samples were correctly detected as misclassifications (Fig. 4).

### Performance of between-dataset classification

The accuracy of cross-dataset predictions depended on the combination of source and target datasets. We first considered the effect of image quality. When the L_L_ images were used to train the NN and then to predict the L_H_ images, the accuracy remained largely constant at 71% for 250 images (Fig. 2b). The accuracy reduction (Δ*Acc*), *i.e*. the reduction in success of predictions compared to the predictions expected from within-dataset classification, also rapidly increased with the number of images, indicating that the training with L_L_ images did not improve the prediction of the L_H_ images (Fig. 5).

**Figure 5.**
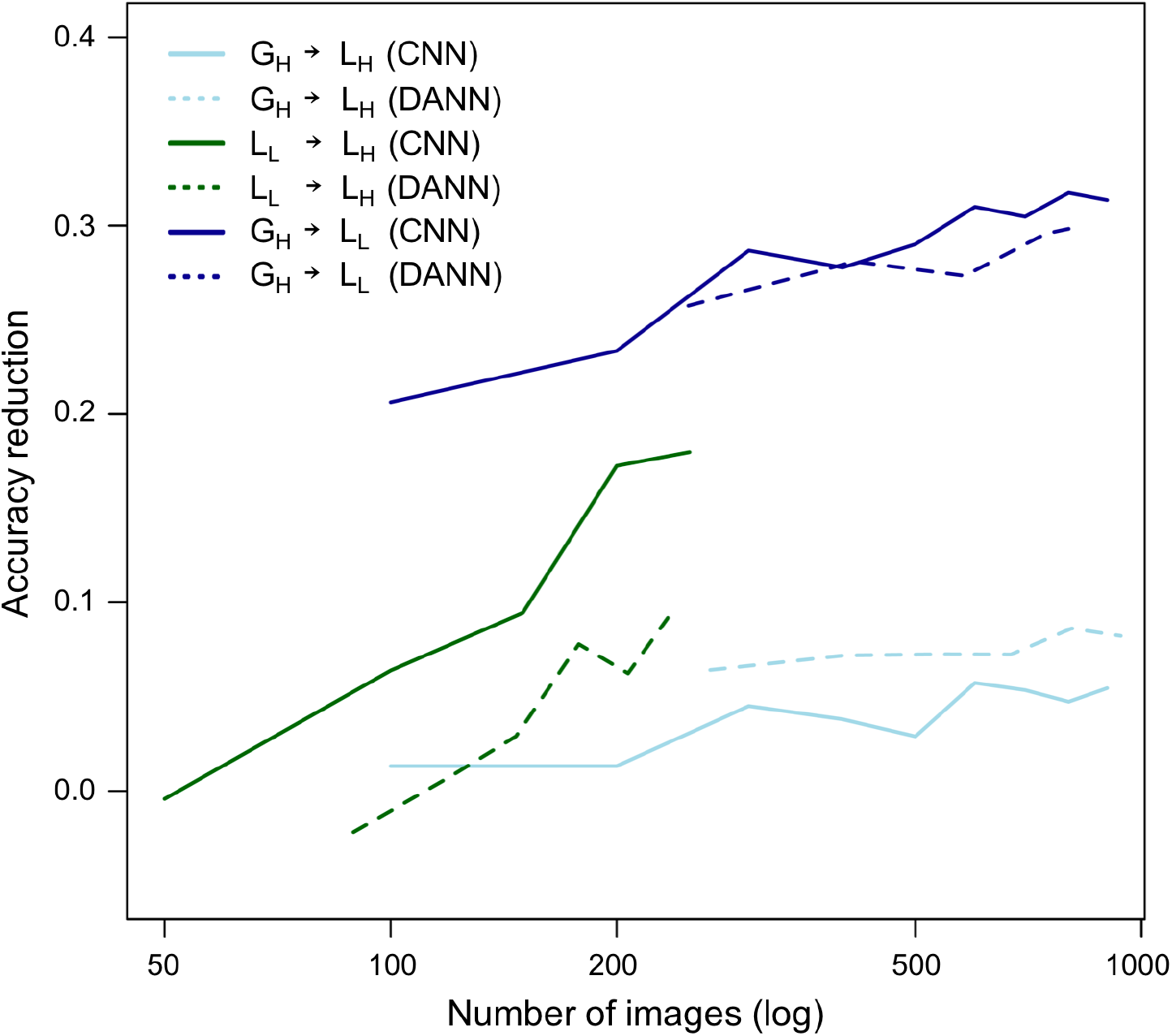
The effect of increasing numbers of images on the accuracy reduction in across-dataset predictions. Subsets of randomly selected images of one dataset are used for training and predicting the class of another set, as indicated by different colours. The x-axis representing the number of images is on a logarithmic scale. Higher accuracy reduction indicates a worse performance on prediction compared to the within-dataset prediction accuracy. The solid and dashed lines represent results of the convolutional neural network (CNN) and domain adversarial neural network (DANN), respectively. Note that only the L_L_ to L_H_ prediction accuracy improved with the use of DANN.

Second, we considered the power of the global dataset to predict the local data, using the G_H_ and the L_H_ as a source-target pair. The prediction accuracy for this comparison was close to the within-G_H_ predictions, with the average accuracy being 79% and the maximum 82% with 900 images (Fig. 2c), indicating that the local set from the Cyprus collection (L_H_) behaves in a similar way as the other local sets contributing to the G_H_ dataset. The accuracy reduction from G_H_ to L_H_ was on average 0.04 and remained almost constant after 300 images (Fig. 5). The power of the G_H_ dataset required the high image quality exhibited by the target (L_H_); when the G_H_-trained model was used to predict the L_L_ images, the accuracy was significantly lower (Fig. 2c). This was also evident from the increased accuracy reduction with increased number of images; whereas the G_H_ to G_H_ predictions improved with more images, the G_H_ to L_L_ predictions did not, resulting in a higher Δ*Acc* (Fig. 5). The dataset classification errors (ε_souce-target_) were 0.20 (G_H_→L_H_), 0.06 (G_H_→L_L_) and 0.01 (L_H_→L_L_), indicating high similarity between the G_H_ and L_H_ images and the distinctiveness of the L_L_.

Table S4 shows a confusion matrix of the G_H_→L_H_ prediction trained by 800 images. Chrysomelidae, Curculionidae and Staphylinidae had recall rates >0.90 (Fig. 3b), but more taxa were incorrectly classified than in the case of the L_H_→L_H_ prediction. No image of Leiodidae and Scaphidiinae, with the available training images <50, was predicted correctly (Fig. 3b). Misclassification mostly affected morphologically similar taxa, *e.g*. the reciprocal confusion of Brentidae and Curculionidae (Table S4).

There was a strong positive correlation between class-wise recall rates and the number of images in the source dataset (rho = 0.77, Fig. 3b). Three taxa with >300 images had recall rates >0.95, while the taxa with <40 images had recall rates <0.4 (Fig. 3b). The effect of the number images on the class-wise precision and F1-score was also positive, but slightly weaker (rho = 0.16 and 0.42, respectively; Fig. 3d,f). Surprisingly, the F1-scores were greatly reduced relative to the recall score for the Chrysomelidae, indicating the precision of the prediction was low even if the recall was high (Fig. 3d,f), *i.e*. the true Chrysomelidae were correctly classified, but many other taxa were incorrectly classified as Chrysomelidae.

As observed in the within-dataset classification, average prediction probabilities of successful predictions (0.98) were consistently higher than the failed predictions (0.84) and *out-of-distribution* samples (0.77). However, failed predictions more frequently had probabilities > 0.95 (Fig. 4b).

#### The performance of the domain adversarial training

The DANN significantly improved the target accuracy of the L_L_ → L_H_ prediction, which involves images from very different setups (Fig. 6). A linear regression model showed that the target accuracy increased by 6.2% (Fig. 6) and the accuracy reduction decreased by 0.060 when the DANN model was used with labeled L_L_ and unlabeled L_H_ images (Fig. 5). The average target accuracy was 79% with 200 labeled L_L_ images and 400 unlabeled L_H_ images (Fig. 6), approaching the same level of accuracy as G_H_→L_H_ predictions.

**Figure 6.**
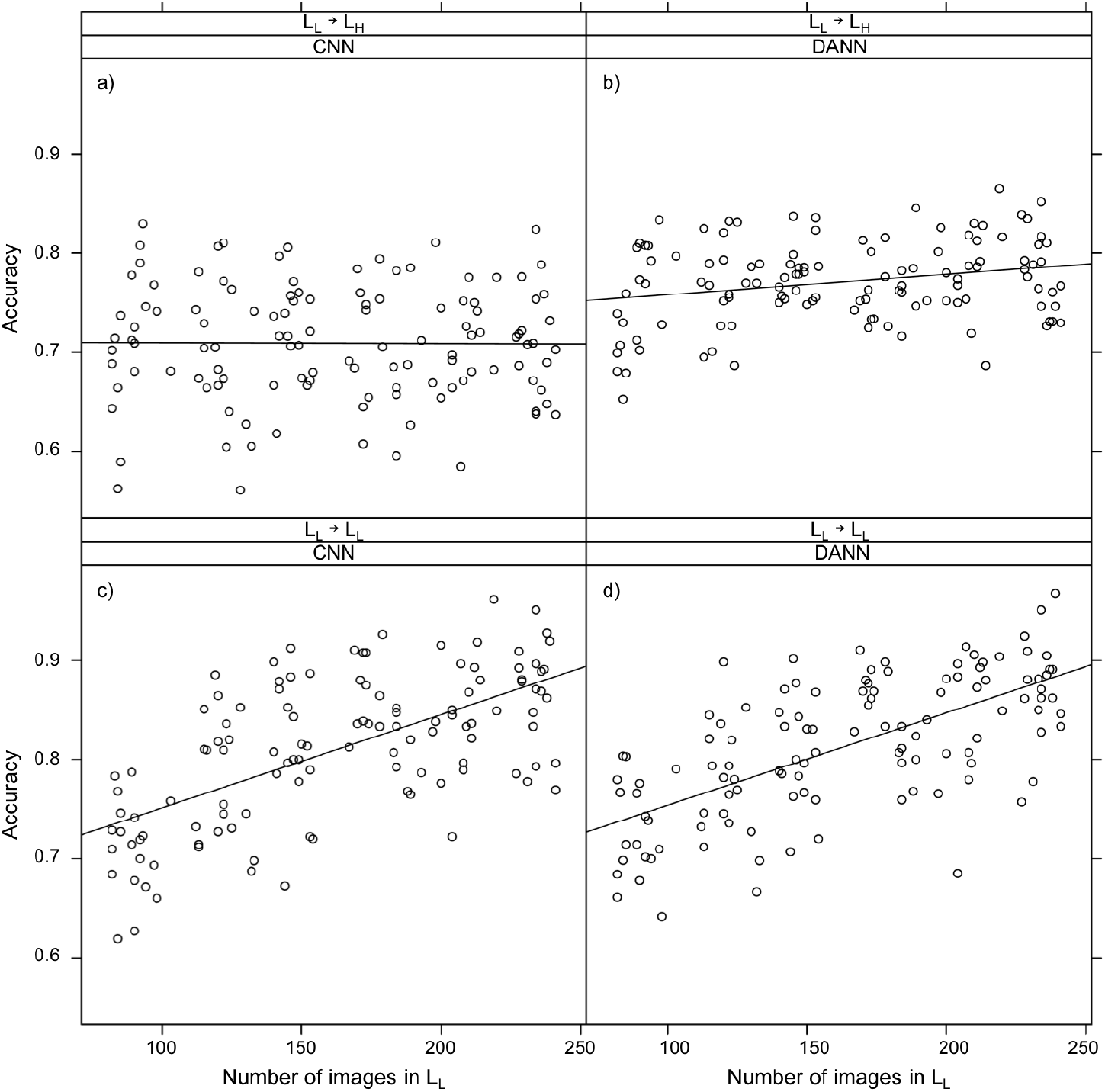
Effect of the number of images on prediction accuracy of the convolutional neural network (CNN, panels a and c) and the domain adversarial neural network (DANN, panels b and d) training for the Local Low Quality (L_L_) and Local High Quality (L_H_) images. Top panels (a and b) represent between-dataset predictions (L_L_→L_H_) while bottom panels (c and d) indicate within-dataset predictions (L_L_→L). Solid lines represent regression lines between the number of images and accuracy. For both between- and within-dataset predictions, models using DANN were trained with a mixed set of randomly selected images from the L_L_ and L_H_ datasets. For other dataset comparisons, see Figures S3 and S4.

On the contrary, the DANN did not improve the target accuracy when the G_H_ was used as a source dataset (Fig. S3 and S4). The G_H_→L_H_ target accuracy was on average 0.75 with 940 labeled G_H_ images and 460 unlabeled L_H_ images (in total 1400 images), which was significantly lower than the between-dataset predictions by the plain NN model (Fig. S3). In the G_H_→L_L_ prediction, a similar trend was observed (Fig. S4) and the target accuracy was not significantly different from the NN.

## DISCUSSION

This work adds to the growing number of studies demonstrating the power of CNNs in image-based taxonomic classification. Specifically, we tested the possibility of classifying specimens from bulk samples of beetles, whereby unknown local samples were classified using a model trained on similarly photographed bulk samples from a global sampling effort. We envision that mixed trap samples in future will be routinely photographed with high-resolution cameras, producing huge numbers of valuable images, but unlike most existing studies that use pinned or cardboard-glued specimens, these images present specimens in diverse angles, habitus, magnification, and lighting (Schneider *et al*. 2021; Wührl *et al*. 2021). We show that these images provide sufficient information for specimens to be identified as members of particular families of Coleoptera. Within a local dataset, classification accuracy regularly reached 95% or more, which is similar to findings from other studies using more standardised photographs from museum collections (*e.g*. ~92% and 96% for Diptera and Coleoptera, respectively; Valan *et al*. 2019). We also confirm that classification performance depends on the number of images used for training (Figs. 2–3), as widely seen in image recognition applications generally (Donahue *et al*. 2013) and in insect classification in particular (*e.g*. >90% recall rates were obtained for taxa with >50 images; Valan *et al*. 2019, 2021). We find that the prediction accuracy generally does not increase further after about 200 images in each of the three datasets used here. However, the degree of accuracy is greatly affected by the image quality and the complexity of the dataset: both the L_L_ (low image quality) and in particular the G_H_ (high complexity) datasets show comparatively low accuracy of predictions if trained on themselves.

### Utility of global databases for classifying local faunas

The critical question in this study is about the success of domain transfer in a situation where the source and target data are from different faunas. We here used the challenging case of the below-ground fauna of a Mediterranean island as the domain target for images trained on a set of above-ground samples from several tropical forests across the globe (the G_H_ set), which presumably do not share any species or genera. However, most local bulk samples, even from such disparate ecosystems, share a similar set at the family level, especially for a small number of species-rich families which are found in similar relative proportions in most samples. We find that the G_H_→L_H_ prediction suffered only low accuracy reduction, confirming the possibility of classifying the high-throughput images from Cyprus by training a convolutional neural network (CNN) model with the global images even though the target species are not present in the training data. We note that the G_H_ dataset is a complex composite of samples from 11 different sites around the globe, collected using a range of different trapping methods (which explains why the within-dataset accuracy was lower than in the other datasets). We argue that this is not necessarily a negative feature, as such a complexity may allow the CNN model trained on this set to capture general family traits of the global fauna and thus make it suitable for a greater range of classification tasks at local level. The high accuracy obtained using the global training set indicates that it is not strictly necessary to create local reference databases for training, when targeting higher taxonomic levels. This finding opens the way for local biodiversity assessment studies around the globe using a universal training set. Global databases have the additional advantage of offering high numbers of images per taxon, which is more difficult to obtain locally, while it is critical for increasing the performance of the CNN-based classification (Fig. 2; Donahue *et al*. 2013; Valan *et al*. 2019, 2021).

Despite the high prediction accuracy of the dataset as a whole, some taxa may show consistently lower classification performance. The primary factor affecting recall and precision is the number of images per family. The required quantity was available only for the largest families (which were also available for the greatest number of countries globally, as a measure of complexity of the training set). However, a few taxa, including the widely sampled Chrysomelidae, showed low F1-scores even with a large number of images (Fig. 3b,d,f). This example is particularly striking because of the high recall rate, but low precision, *i.e*. while most specimens of Chrysomelidae in the sample are identified with a high prediction probability, the model misclassifies a lot of them and incorrectly assigns specimens of other families to them. The Chrysomelidae behaves poorly against the local L_H_ model, but this is commensurate with a low representation of images (Fig. 3a,c,e). The finding may suggest a negative impact of within-family morphological disparity on classification precision, possibly only present in the wider G_H_ dataset. Interestingly, Chrysomelidae also showed low classification performance in the study of Valan *et al*. (2019). The family is composed of morphologically rather distinct subfamilies, and an increased number of images may help to unveil the subclasses generating low performance models.

### Lessons from combining DANN with differing databases

We show that photographs taken from similar imaging setups (G_H_ and L_H_) are readily used for between-region image classifications while images taken by a conventional stereoscope (L_L_) exhibited a large accuracy reduction for the prediction of the local high quality dataset. Considering the nearly identical taxonomic composition of the L_H_ and L_L_ datasets, the large accuracy reduction indicates a negative impact of the original image quality and the lack of standardization between the target-source pairs. The overall dissimilarity of L_H_ from G_H_ and L_L_ measured by dataset classification errors also suggest a negative effect of non-standardized imaging on prediction performance. These results are in accordance with the reduction in classification accuracy observed by other studies comparing different imaging procedures, *e.g*. training with high-resolution museum specimens to predict field images (Knyshov *et al*. 2021). The application of alternative algorithms may overcome limitations resulting from the usage of highly different images taken by unstandardized imaging conditions. In the current study, we could successfully ameliorate the accuracy reductions between L_H_ and L_L_ using DANN, a method designed for domain adaptation (Ganin *et al*. 2016). However, in other combinations of datasets such as G_H_ and L_H_, the DANN did not improve the target prediction performance. This may be due to poor hyperparameter tuning or insufficient training of the model with a complex loss function (Kouw & Loog 2021). Nevertheless, our study would offer some evidence that DANN (or domain adaptation techniques in general) can be considered a method of choice when a standardized image acquisition is not available.

### Improvements from using alternative metrics for model performance

While CNN-based image classification for biodiversity assessment is becoming increasingly popular, its performance is not always assessed with a broad set of performance metrics. As observed in Chrysomelidae, the reduction of performance was only detectable in the multiclass precisions and F1 scores, but not in the recalls, which revealed a specific difficulty in the classification of this group. Given the inferential power of these performance metrics, we encourage their integration in biodiversity-related applications.

Another overlooked metric is the confidence of predictions. We could detect failed predictions and potential *out-of-distribution* samples by setting a threshold value on the probabilities. In accordance with Hendrycs & Gimpel (2017), such misclassified or *out-of-distribution* samples were predicted with consistently lower prediction probabilities. Because *out-of-distribution* samples are common in biodiversity surveys, detection of unknown target samples based on low prediction confidence is particularly useful. A potential difficulty of this approach is that calibration of the threshold requires extra data. Conventional deep neural networks can be uncalibrated, that is, prediction probabilities do not precisely reflect prediction accuracy (Guo *et al*. 2017). Such uncalibrated models can make an incorrect prediction with excessively high confidence. This overconfident failure is noticeable in our analysis (Fig. 4b). Therefore, additional labeled samples are required to set a robust threshold for the identification of failure and *out-of-distribution* samples. Methods for explicit calibration of prediction probabilities or detection of *out-of-distribution* samples without additional data (*e.g*. Hsu *et al*. 2020; Mukhoti *et al*. 2020) are being actively developed in the machine learning field, and applying those methods is a potential future direction. As the DANN could remove the dataset biases caused by the imaging instruments, the purpose-specific models will expand the possibility of machine learning applications to biodiversity surveys (see Høye *et al*. 2020).

### Building the global database for CNN-based classification

As new images become available for ever more species, the reference library for taxonomic identification is rapidly growing. Our training set was derived from various biodiversity hotspots around the world and was classified at the level of families only. Given the geographic and taxonomic distance of these samples, the family category is the only meaningful level exhibiting overlap of source and target, but conceivably the methodology could be applied at lower levels, *e.g*. genera, if more similar samples had been used. The current set of images is limited with regard to the number of families (classes) and number of images per family (intra-class variability), resulting in *out-of-distribution* errors and prediction errors, respectively. Both issues can be addressed with a wider selection of images, *e.g*. those available from the SITE-100 project (*https://www.site100.org*) taken with similar equipment. Based on our results, any future image collection should consider the need for standardisation, including that imaging should use the same aspect, *e.g*. dorsal view for Coleoptera (also see Hansen *et al*. 2019), uniform background across images (preferably a clear colour without textures), clear separation of specimens in the photographs, and similar optical equipment and magnification. The exact parameters remain to be explored within and across studies, but standardisation of imaging is critical to transferability when rolling out large-scale efforts for image-based classification in biodiversity studies. As part of this effort, image segmentation should be improved and automated (Schneider *et al*. 2021; Schwartz & Alfaro 2021), to increase our capability for rapidly generating ‘clean’ and individual-based image databases extracted from bulk samples. A potential bottleneck is the need to expand the training set gradually, which generally requires recomputation of the model when new classes are added, although recent update methods may simplify this process (Hadsell *et al*. 2020). A second issue affecting the accuracy of predictions is the “category bias” from inconsistent categorisation and labeling of the training set itself. In the current study, images in the training set were classified from the images by recognising the overall gestalt of a family. These class labels were straightforward for most groups, but identification of some beetle families may be compromised due to images that obscured appendages or other key traits, especially in small-bodied Leiodidae, Latridiidae or Cryptophagidae, which may have contributed to the prediction errors seen in these families (Table S3). Thus, corrections to the class labels in the database may be required, possibly by integrating image-based classification with widely-established DNA barcoding and phylogenetic placement methods that confirm the class membership. Likewise, combining image acquisition for biodiversity assessment with metabarcoding could be instrumental for validating or improving genetic-based inferences (Yang *et al*. 2021) or estimating biomass and abundance (*e.g*. Høye *et al*. 2020; Schneider *et al*. 2021). Metabarcoding studies often lose morphological information of specimens, but imaging could be accommodated as a routine step before the DNA extraction of bulk samples.

## CONCLUSIONS

To our knowledge, this is the first attempt of domain transfer for taxonomic classification of an entirely unknown dataset, as a key element of using image-based identification in biodiversity studies at the global scale. We show that the approach is highly feasible, but needs careful consideration of the imaging procedure, the algorithmic approach, and the choice of training sets. The future vision of this approach is an increasingly complete set of images, covering the diversity of major taxonomic groups, against which samples from any ecosystem and biogeographic region can be classified at a certain hierarchical level (*e.g*. families of beetles). In our approach we lack the close alignment of source and target that would guarantee high transferability, but at the expense of lower generalization capability. Further studies are required to study the trade-offs and to establish best practice for the specific research question at hand. Once a stable expanded image database has been created, it can be used for broader applications in biodiversity research and monitoring, potentially building a global model applicable to any sampling site and possibly used while still in the field.

## DATA AVAILABILITY STATEMENT

The code to reproduce the analysis and image databases will available for download from Dryad Digital Repository, upon acceptance.

## ACKNOWLEDGMENTS

This work was supported by the iBioGen project, which has received funding from the European Union’s Horizon 2020 research and innovation programme under grant agreement No. 810729. We are grateful to Richard Turney and Thomas J. Creedy (Natural History Museum, London) for advice and support during bulk-sample imaging, and Takashi Imai (Shiga University) for helpful advice on deep learning methods. We would like to extend our gratitude to Andreas Dimitriou for help during sample imaging, and Konstantinos Ntatsopoulos for support in the taxonomy of Cyprus beetles. V.N. was supported by a postdoctoral contract under the iBioGen project and a “*Juan de la Cierva-Formación’’* postdoctoral fellowship (grant: FJC2018-035611-I) funded by MCIN/AEI/10.13039/501100011033. T.F. was supported by JSPS KAKENHI (grant number: 20K06824).

## AUTHOR’S CONTRIBUTIONS

T.F., V.N., A.P. and A.P.V. conceived and led the study. V.N and E.M performed fieldwork for Cyprus data and generated the associated image dataset. T.F., V.N. and A.P. conceived the methodological approach and T.F. performed all deep learning analyses. T.F., V.N., A.P. and A.P.V. wrote the manuscript. All authors contributed critically to the draft and gave final approval for publication.

## SUPPLEMENTARY MATERIAL

**Figure S1.**
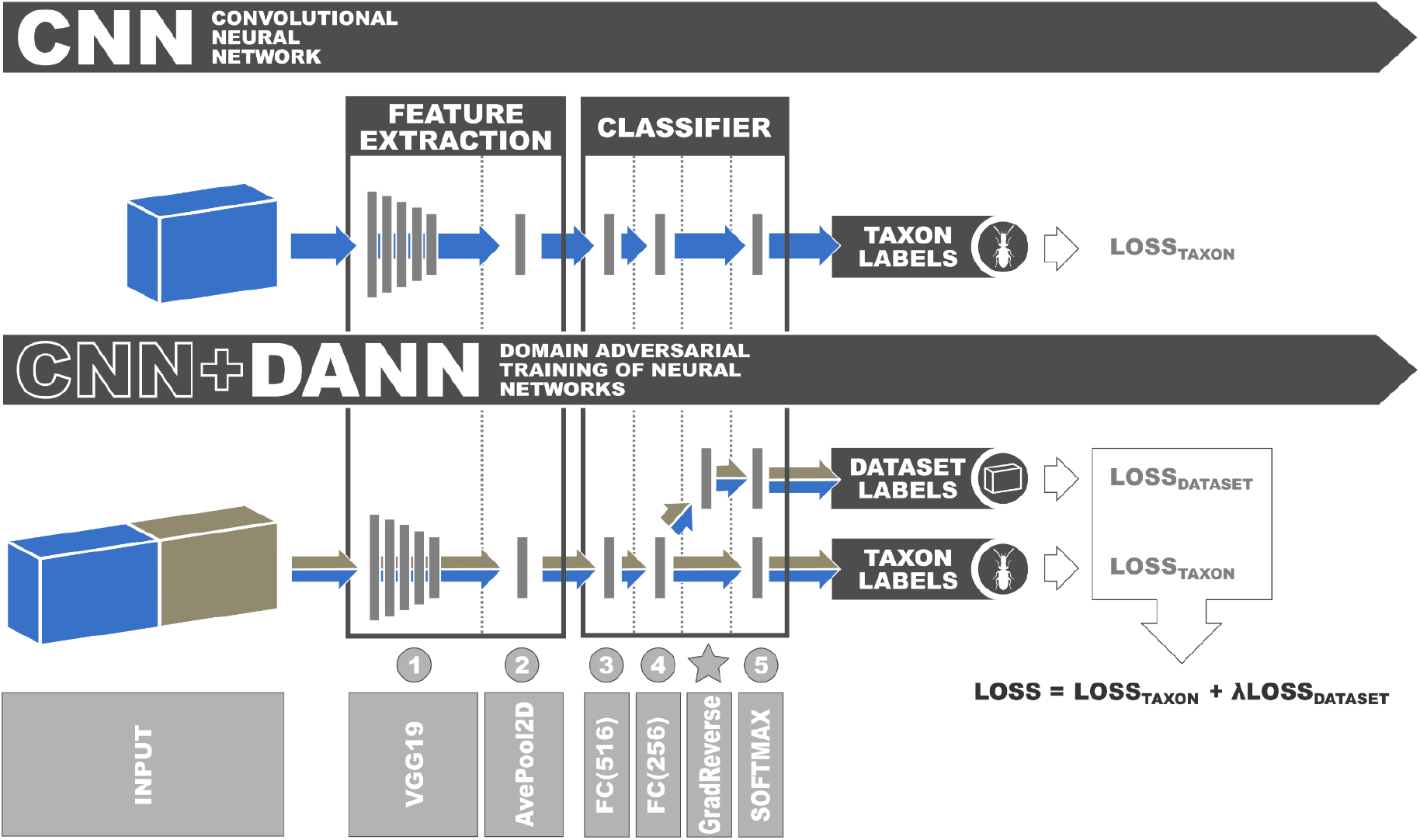
Detailed scheme of architectures of the convolutional neural network (CNN) and domain adversarial neural networks (DANN) used in this study. For DANN, the outputs of the second fully connected (FC256) layer were forwarded to two softmax layers for taxon and dataset classification. A gradient reversal layer was inserted between the FC(256) and dataset softmax layers to achieve domain adversarial training as proposed in Ganin *et al*. (2016).

**Figure S2.**
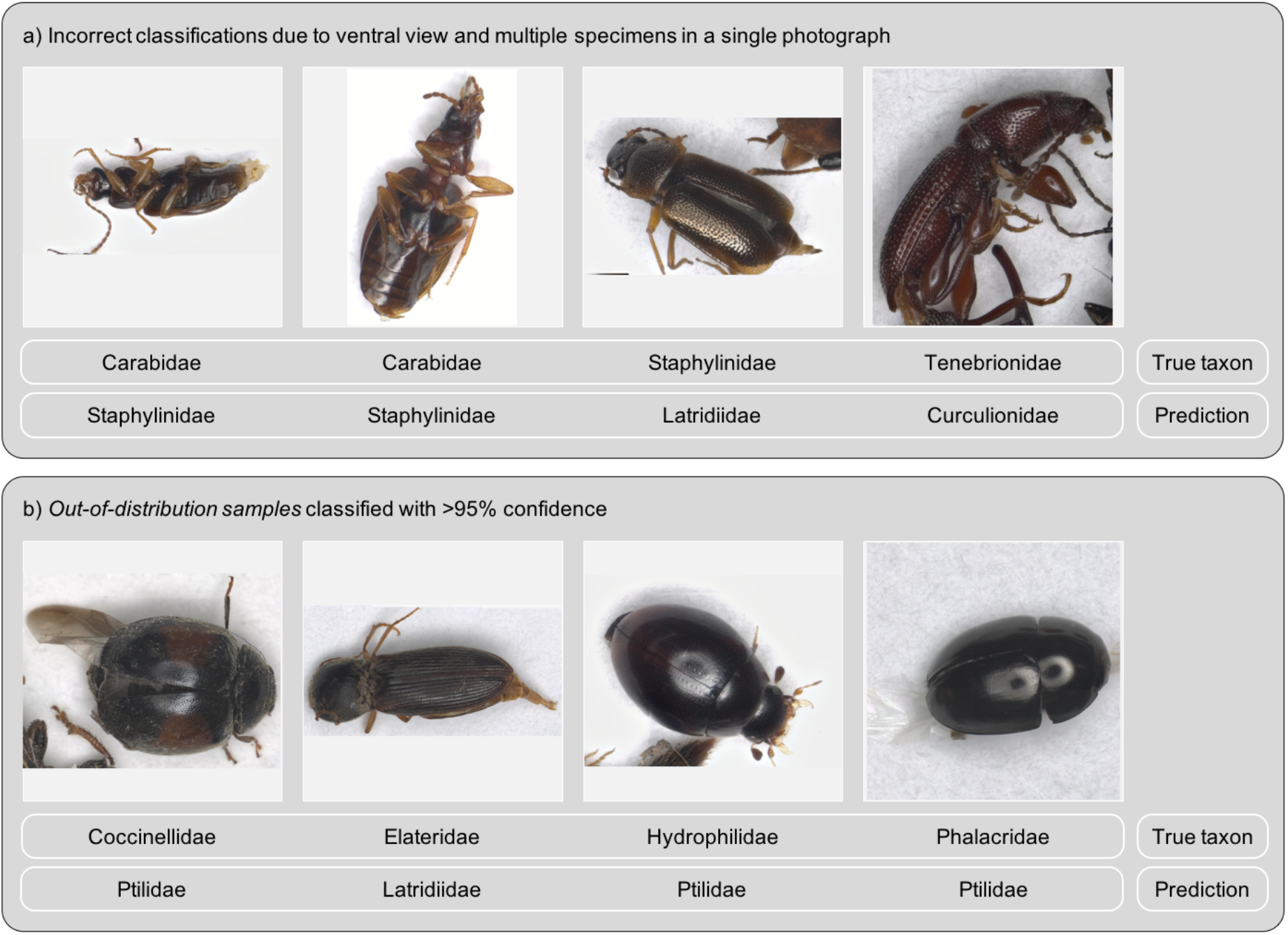
Exemplar images of incorrect classification.

**Figure S3.**
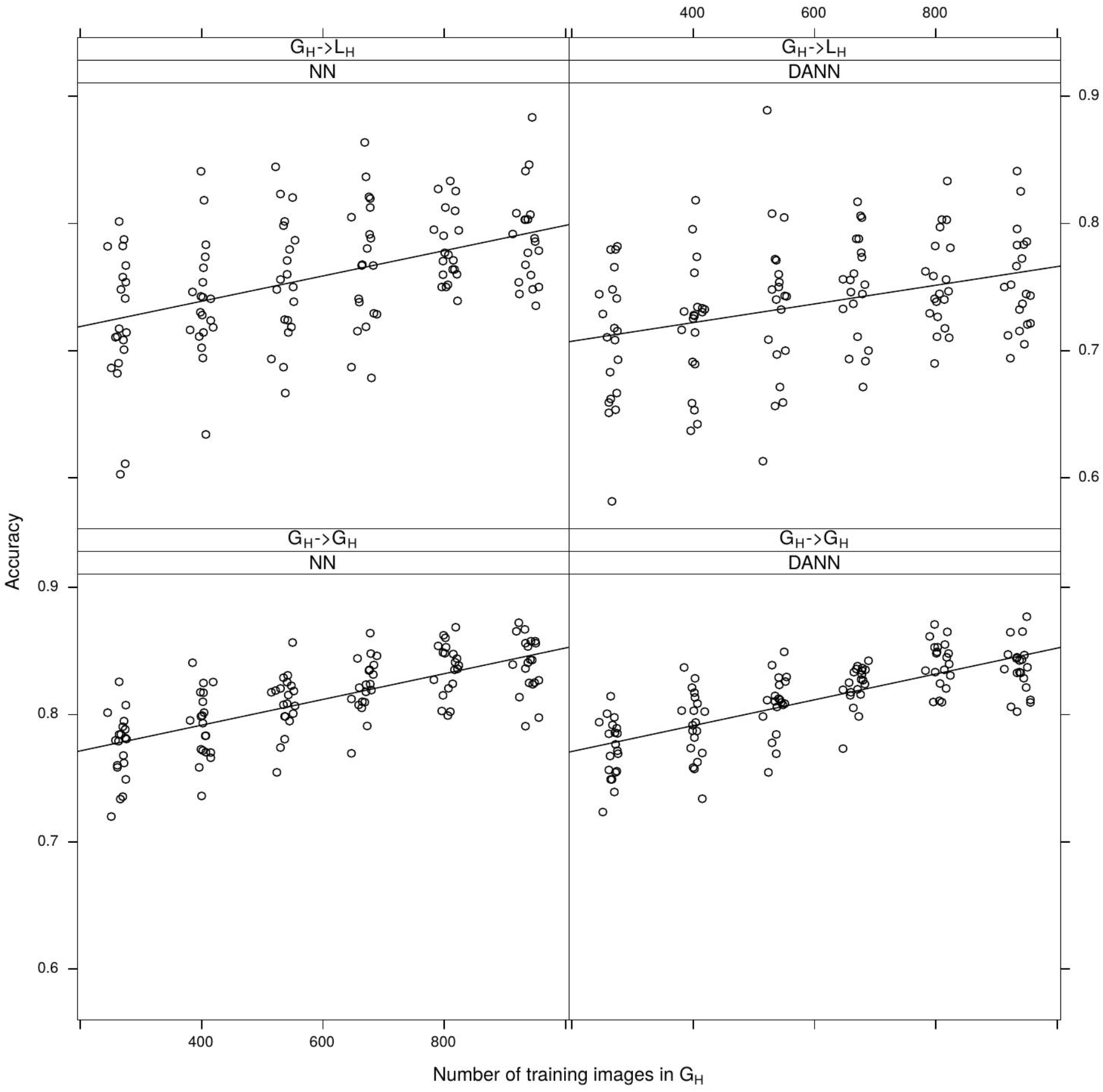
Effect of the number of images on prediction accuracy of the convolutional neural network (CNN, left panels) and the domain adversarial neural network (DANN, right panels) training for the *Local High Quality* (L_H_) and *Global High Quality* (G_H_) images. G_H_→L_H_ (top panels) indicates between-dataset predictions, while G_H_→GH (bottom panels) represent within-dataset predictions. Solid lines represent regression lines between the number of images and accuracy. For both between- and within-dataset predictions, models using DANN were trained with a mixed set of randomly selected images from the G_H_ and L_H_ datasets.

**Figure S4.**
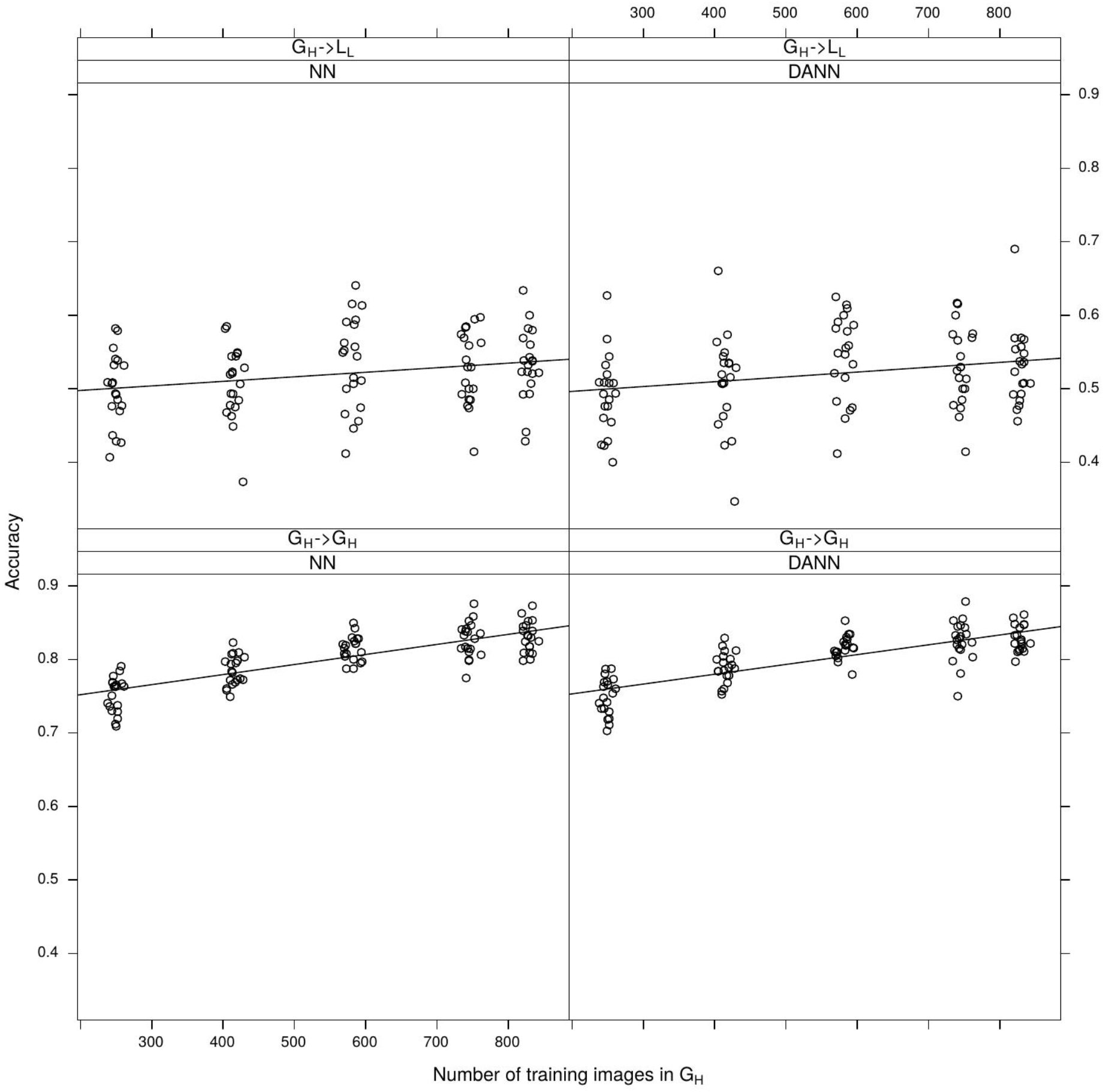
Effects of the number of images on prediction accuracy of the convolutional neural network (CNN, left panels) and the domain adversarial neural network (DANN, right panels) training for the *Local Low Quality* (L_L_) and *Global High Quality* (G_H_) images. G_H_→L_L_ (top panels) indicates between-dataset predictions, while G_H_→GH (bottom panels) represent within-dataset predictions. Solid lines represent regression lines between the number of images and accuracy. For both between- and within-dataset predictions, models using DANN were trained with a mixed set of randomly selected images from the G_H_ and L_L_ datasets.

**Table S1.**
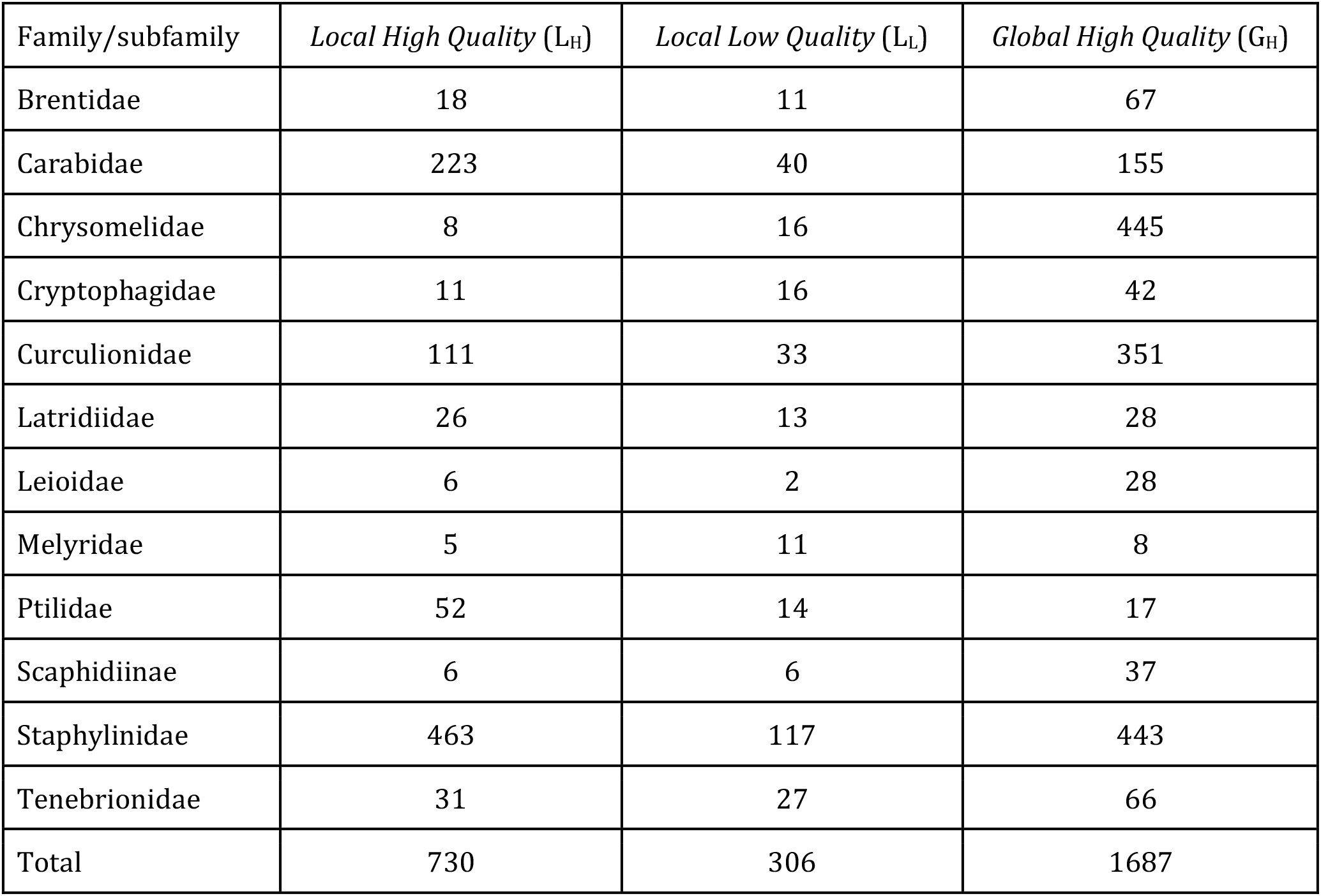
Number of images per taxa and dataset used in the present study.

**Table S2.** Sampling sites for the *Global High Quality* (G_H_) dataset. Images were obtained from a global catalogue of Coleoptera specimens available at *https://www.flickr.com/photos/site-100/*

| Country | Locality | Number of images | Latitude | Longitude | Elevation (m) |
| --- | --- | --- | --- | --- | --- |
| Borneo | Poring, Sabah | 151 | 6° 03' N | 116° 42' E | 550 |
| Ecuador | Yasuni, Onkone Gare | 829 | 0° 39' 30.05" S | 76° 27' 9.56" W | 216 |
| Equatorial Guinea | Djibloho, Oyala | 193 | 1° 36' 27.40" - 1° 37' 13.10" N | 10° 51' 26.60" - 10° 52' 52.00" E | 670 |
| French Guiana | Nouragues | 161 | 4° 05' 15.70" N | 52° 40' 52.70" W | 130 |
| Honduras | Cortes | 228 | 14° 50' 15.00" N | 87° 53' 43.00" W | 415 |
| México | Chamela | 132 | 19° 29' 54.70" N | 105 °02' 38.40" W | 97 |
| India | Mizoram | 46 | 23° 27' 00.00" N - 23° 47' 24.00" N | 92° 36' - 92° 45' E | 800-1600 |
| Panamá | Cerro Hoya | 306 | 7° 14' N | 80° 53' W | 550 |
| Panamá | Santa Fe | 307 | 8° 32' N | 81° 07' W | 550 |
| South Africa | Hogsback | 60 | 32° 36' 21.60" S | 26° 57' 43.30" E | 1070 |

**Table S3.** A scaled confusion matrix for the prediction of *Local High Quality* (L_H_) images based on a training set of 400 randomly selected images from the same dataset (L_H_). Rows represent the predicted classes and columns the true classes. The matrix was scaled so that each column sums up to one.

|  | Brentidae | Carabidae | Chrysomelidae | Cryptophagidae | Curculionidae | Latridiidae | Leiodidae | Melyridae | Ptiliidae | Scaphidiinae | Staphylinidae | Tenebrionidae |
| --- | --- | --- | --- | --- | --- | --- | --- | --- | --- | --- | --- | --- |
| Brentidae | 0.757 |  |  |  |  |  |  |  |  |  |  |  |
| Carabidae |  | 0.952 |  |  |  |  |  |  |  | 0.429 | 0.003 |  |
| Chrysomelidae |  |  | 0.154 |  |  |  | 0.231 |  |  |  |  |  |
| Cryptophagidae |  |  | 0.385 | 0.471 |  |  |  |  |  |  |  |  |
| Curculionidae | 0.135 |  |  |  | 0.977 | 0.098 |  | 0.250 |  |  | 0.010 | 0.213 |
| Latridiidae |  |  |  | 0.235 | 0.023 | 0.820 |  | 0.500 |  |  | 0.003 | 0.138 |
| Leiodidae |  |  | 0.462 |  |  |  | 0.769 |  |  |  |  |  |
| Melyridae |  |  |  |  |  |  |  | 0.000 |  |  |  |  |
| Ptiliidae |  |  |  | 0.176 |  |  |  |  | 1.000 |  |  |  |
| Scaphidiinae |  |  |  |  |  |  |  |  |  | 0.571 |  |  |
| Staphylinidae | 0.108 | 0.048 |  | 0.118 |  |  |  | 0.250 |  |  | 0.983 | 0.038 |
| Tenebrionidae |  |  |  |  |  | 0.082 |  |  |  |  |  | 0.613 |

**Table S4.** A scaled confusion matrix for the prediction of *Local High Quality* (L_H_) images based on a training set of 800 randomly selected *Global High Quality* (G_H_) images. Rows represent the predicted classes and columns the true classes. The matrix was scaled so that each column sums up to one.

|  | Brentidae | Carabidae | Chrysomelidae | Cryptophagidae | Curculionidae | Latridiidae | Leiodidae | Melyridae | Ptiliidae | Scaphidiinae | Staphylinidae | Tenebrionidae |
| --- | --- | --- | --- | --- | --- | --- | --- | --- | --- | --- | --- | --- |
| Brentidae | 0.444 |  |  |  | 0.072 |  |  |  |  |  |  |  |
| Carabidae |  | 0.655 |  | 0.091 |  | 0.038 |  |  |  | 0.167 |  | 0.065 |
| Chrysomelidae |  | 0.103 | 1.00 | 0.273 |  | 0.154 | 0.667 |  | 0.231 | 0.667 | 0.013 | 0.129 |
| Cryptophagidae |  |  |  | 0.273 |  | 0.038 |  |  | 0.019 |  |  |  |
| Curculionidae | 0.444 | 0.004 |  |  | 0.901 | 0.577 |  | 0.80 | 0.250 |  | 0.004 | 0.548 |
| Latridiidae |  |  |  | 0.091 |  | 0.038 |  |  |  |  |  |  |
| Leiodidae |  |  |  | 0.091 |  |  | 0.000 |  | 0.077 |  |  |  |
| Melyridae | 0.056 |  |  |  |  |  |  | 0.20 |  |  |  |  |
| Ptiliidae |  |  |  | 0.091 |  |  |  |  | 0.365 |  |  |  |
| Scaphidiinae |  | 0.004 |  |  |  |  |  |  |  | 0.000 | 0.004 |  |
| Staphylinidae |  | 0.233 |  | 0.091 | 0.027 |  | 0.333 |  | 0.058 |  | 0.978 | 0.097 |
| Tenebrionidae | 0.056 |  |  |  |  | 0.154 |  |  |  | 0.167 |  | 0.161 |

## Notes

### Competing Interest Statement

The authors have declared no competing interest.

